# Emap2sec+: Detecting Protein and DNA/RNA Structures in Cryo-EM Maps of Intermediate Resolution Using Deep Learning

**DOI:** 10.1101/2020.08.22.262675

**Authors:** Xiao Wang, Eman Alnabati, Tunde W. Aderinwale, Sai Raghavendra Maddhuri Venkata Subramaniya, Genki Terashi, Daisuke Kihara

## Abstract

An increasing number of density maps of macromolecular structures, including proteins and protein and DNA/RNA complexes, have been determined by cryo-electron microscopy (cryo-EM). Although lately maps at a near-atomic resolution are routinely reported, there are still substantial fractions of maps determined at intermediate or low resolutions, where extracting structure information is not trivial. Here, we report a new computational method, Emap2sec+, which identifies DNA or RNA as well as the secondary structures of proteins in cryo-EM maps of 5 to 10 Å resolution. Emap2sec+ employs the deep Residual convolutional neural network. Emap2sec+ assigns structural labels with associated probabilities at each voxel in a cryo-EM map, which will help structure modeling in an EM map. Emap2sec+ showed stable and high assignment accuracy for nucleotides in low resolution maps and improved performance for protein secondary structure assignments than its earlier version when tested on simulated and experimental maps.

## Introduction

Recent years have witnessed rapid advances in structural determination of biological molecules using cryo-electron microscopy (cryo-EM)^1,2^. The number of determined cryo-EM maps deposited in the public database, EMDB^3^, is growing exponentially; and moreover, the fraction of high resolution (e.g. better than 4 Å) maps among them shows a steady increase. Despite the remarkable progress of cryo-EM, there are still a substantial fraction of maps determined at intermediate or low resolutions. Different factors control the achieved resolution of EM maps including conformational or compositional heterogeneity of EM samples, noise level introduced by low electron doses, and inaccurate alignment of the two dimensional particle images^4^.

Depending on the resolution of an EM map, structural information that can be extracted from the map and computational tools appropriate for the task will apparently differ^5^. For maps at a near atomic resolution (better than 3 Å) or at the subsequent level of the resolution (∼4 Å), full atom structure models can be usually built using tools for atomic structure modeling^6-8^, de novo main-chain tracing ^9,10^, structure refinement^11,12^ or combinations thereof. As the resolution becomes worse, extracting structure information becomes more challenging. In a map of intermediate resolution (∼4 to 10 Å), protein secondary structure elements can be often detected even in cases where tracing a full sequence is difficult. Tools for this task include those which detect typical local densities that correspond to an α-helix and β-sheet^13,14^. Recent methods benefit from the strong image recognition capabilities of deep learning to extract the local and global density features of EM maps^15,16^. Identified structural fragments in a map can be used as clues for tracing a full-length protein chain, to identify known structures from the database that agree with the fragments^17^, or to identify structural domains of a complex in the map by their secondary structure content.

Previously, we have developed a method named Emap2sec^16^ for detecting protein secondary structure elements in EM maps of intermediate resolutions. Emap2sec showed promising results and provided a novel approach to structural interpretation of maps at intermediate resolution. However, Emap2sec’s detection was limited only to maps of proteins. Here, we have extended the method to detect both protein secondary structure elements and nucleic acids in EM maps of intermediate resolution. The new method, Emap2sec+, uses a more advanced convolutional neural network architecture, Resnet^18^, than its predecessor, Emap2sec, and performs a four-class classification, α-helix, β-strand, coil (other structure type), or DNA/RNA, for each voxel in an EM map. Protein nucleic-acid interactions are the core of many essential biological processes including transcription, translation, cell division, and replication. Despite the importance of investigating structures of protein-nucleic acid interactions, there are not many tools available for DNA/RNA structure modeling. Currently, 1340 EM map entries in EMDB have protein-nucleic acid complexes, which is around 13% of the total number of EM maps at EMDB. Recently, a similar tool, Haruspex^19^, was developed, which detects nucleotides and protein secondary structure in EM maps. However, Haruspex is designed for high resolution EM maps of 4 Å or better; thus the aim of the current tool, Emap2sec+ is very different. Haruspex is not suitable for maps of intermediate resolution by design, as we will show later.

We tested Emap2sec+ on two datasets, a dataset of simulated EM maps from 108 complex structures of proteins and nucleic acids at resolution 6 and 10 Å as well as a dataset of 83 experimental EM maps of a resolution between 5 and 10 Å. Emap2sec+ showed a high accuracy for nucleotide detection while maintaining comparable, if not achieving better, protein secondary structure detection performance to Emap2sec.

### The architecture of Emap2sec+

Emap2sec+ scans a cryo-EM map with a voxel of 11^3^ Å^3^ with a stride of 2 Å and outputs a detected structure at the center of the voxel, which is either DNA/RNA or a protein secondary structure (i.e. α-helices, β-sheets and what we term ‘other structures’). Thus, Emap2sec+ classifies a voxel into four different structural classes. The deep neural network architecture of Emap2sec+ is illustrated in **Figure 1**. Emap2sec+ has two phases: In the first phase, Emap2sec+ performs five independent evaluations for an input voxel (**Fig. 1a**). Among the five channels, four channels perform binary evaluations, where each of them examines whether the voxel contains a particular structure or not. On the other hand, the last channel performs a four-class classification, outputting four probabilities of the four structures (**Fig. 1b**). Then, in the phase 2 network, the eight probability values that are output from the phase 1 network are collected for the query voxel as well as the 7^3^ neighboring voxels. These values are fed into another deep convolutional neural network, which outputs four probability values for the four structures (**Fig. 1b**). The purpose of this network is to consider detected structures of neighboring voxels in making the decision of the query voxel, which consequently has a smoothing effect to the structural assignment in the EM map.

**Figure 1.**
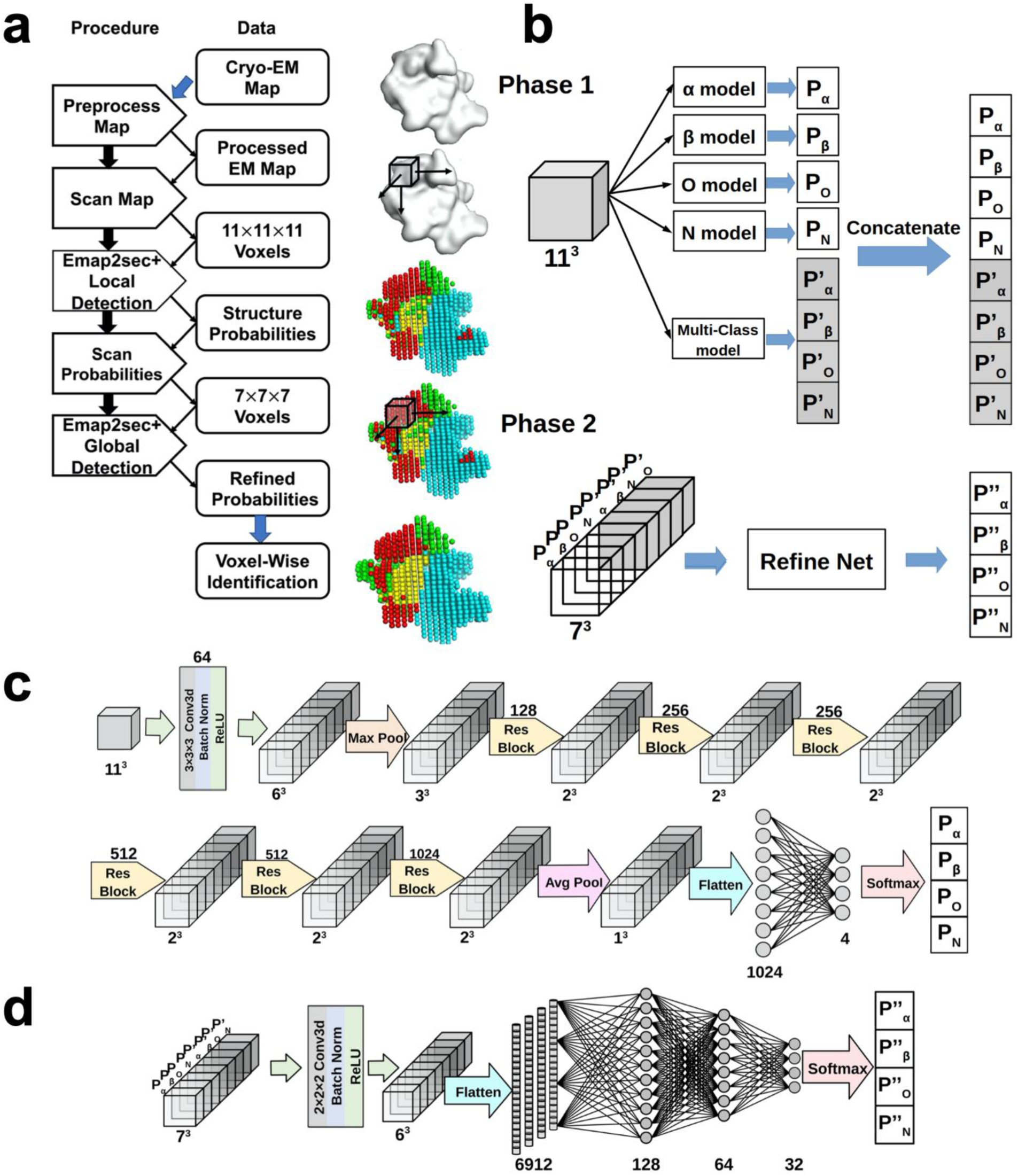
The network Architecture of Emap2sec+. Emap2sec+ scans an EM map with a voxels of 11*11*11 Å^3^ of size and outputs the probabilities that the voxel has α helix, β strand, other structures, or DNA/RNA in the middle of the voxel. It consists of two networks, phase1 and phase 2, where the phase 2 network refines the initial output by considering assignments given to neighboring 7×7×7 voxels by the phase 1 network. **a**, logical steps of the pipeline. **b**, the architecture of the phase 1 and phase 2 networks. The phase 1 consists of 4 binary classifies and one multi (four) -class classifier. The phase 2 network takes outputs from the phase 1 network and outputs refined, final probability values. **c**, a detailed representation of the phase 1. It uses 6 Residual blocks (Supplementary Figure S1). **d**, a detailed architecture of the phase 2. The main part is a fully connected network.

A detailed architecture of the five channels in the phase 1 network is illustrated in **Fig. 1c**. For an input voxel of size 11^3^ Å^3^, a convolutional block is applied, which consists of a convolutional layer with 64 filters of size 3^3^ Å^3^, a 3D batch normalization layer^20^ (the batch size is set to 256), and a ReLU activation, which produces 64 voxels of 6^3^ Å^3^. Then, the voxels are passed to a max pooling layer with size 2^3^ Å^3^. After that, each voxel is connected to 6 3D Residual blocks ^18,21^ (Supplementary Fig. S1) with128, 256, 256, 512, 512, and 1024 filters of size 3^3^ Å^3^. At the last step, an average pooling layer of size 2^3^ Å^3^ is applied resulting in a feature vector of 1024 values, which is connected to a fully connected layer to give the final probability values. The phase 2 network incorporates probability values of 7^3^ neighboring voxels. The input is processed with a convolutional block that consists of a 3D convolutional layer with 32 filters of size of 2, a 3D batch normalization layer, and a ReLU activation layer (the batch size is 256), followed by a fully connected network (**Fig. 1d**). The stride of the convolutional layer was 1. Finally, a softmax operation is applied to the output probability values of the four structure types. The networks were trained on simulated map data for applying to simulated maps while for experimental maps, training was performed with experimental maps (**Supplementary Table S1**). Training details are provided in Methods.

### Structure detection on simulated maps

We investigated the performance of Emap2sec+ on a dataset of 108 non-redundant simulated maps each at 6 and 10 Å as well as on a set of 83 experimental maps. The neural networks were trained separately for simulated and experimental maps using independent training sets. Refer to the Methods section for details of the dataset construction and parameter training of the networks.

The performance of Emap2sec+ was evaluated at three different levels, voxel-based, Q4 (residue/nucleotide-based), and segment-based. In **Figure 2a**, voxel-based F1-score and residue-based accuracy are presented in bar graphs for simulated maps at 6 Å and 10 Å. The segment-based accuracy and other result values are provided in **Supplementary Table S2**. Accuracy values of individual maps of 6 and 10 Å from the phase 1 and 2 networks are made available in **Supplementary Table S3**.

**Figure 2.**
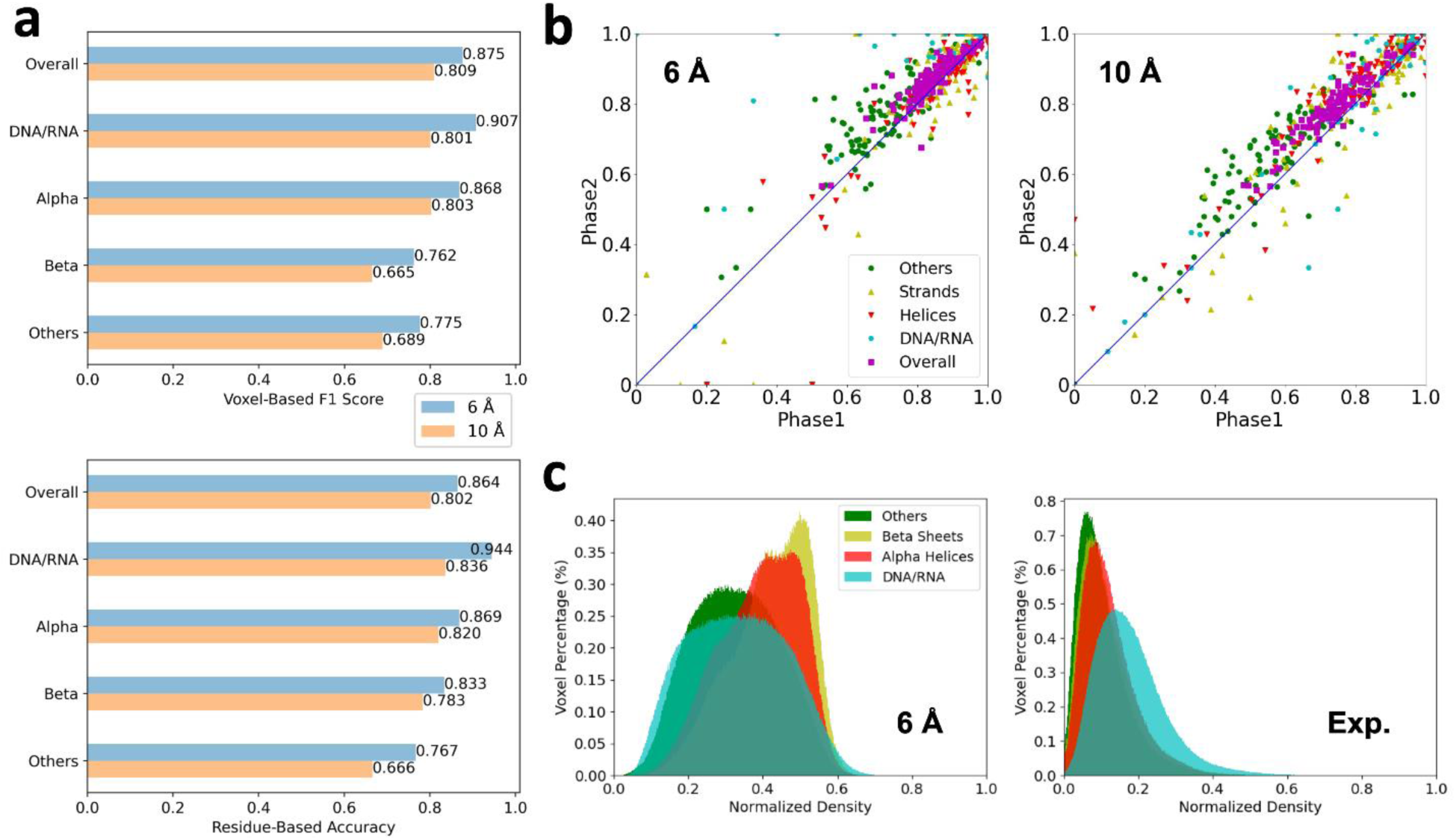
The structure detection performance on the simulated map dataset. The dataset consists of 108 structures computed at 2 different resolutions, 6 Å and 10 Å. **a**, Voxel-based F1 score and Q4 residue/nucleotide-based accuracy for 6 Å (blue) and 10 Å (orange) maps. **b**, Comparison of Q4 of the phase1 and phase2 network outputs for each of 108 test simulated maps computed at 6 Å and 10 Å. Green, other structures; yellow triangles, β strands; red triangles, α helices; cyan, DNA/RNA; magenta, overall Q4. **c**, Distribution of normalized densities of four structural classes in the 108 test maps simulated at 6 Å and experimental maps (Exp.). Corresponding distributions of 108 test simulated maps at 10 Å are shown in **Supplementary Figure S2**.

For the 6 Å maps, Emap2sec+ achieved an overall F1-score of 0.875 and Q4 score of 0.864. Comparing the four structure classes, DNA/RNA had the highest F1-score (0.907) and the Q4 score (0.944). The good performance on DNA/RNA detection was not due to a distinct density from proteins. **Figure 2c** shows that the density distribution of DNA/RNA have significant overlaps with protein structure classes and is not distinctive particularly in simulated maps. In experimental maps, the density of nucleic acids tends to be higher than proteins (**Fig. 2c**, density distributions for 10 Å maps are shown in **Supplementary Figure S2**). Among the three classes in proteins, α helices, β strands, and other structures, α helices were best detected, which is consistent with the previous version of the software, Emap2sec^16^. When only the three protein structure classes are considered, the Q3 score were 0.846 and 0.780 for 6 Å and 10 Å maps, respectively (**Supplementary Table S3)**, which are better than results shown in Table 1 of the Emap2sec paper^16^. Thus, adding the additional class of DNA/RNA in Emap2sec+ did not deteriorate classification for protein secondary structure classes.

Comparing voxel-based accuracy (**Supplementary Table S2**) and Q4, Q4 are higher in all structural classes. Considering that the residue/nucleotide-based assignments are made by a majority vote from voxels, it indicates that neighboring voxels tend to have consistent assignments, which facilitate users to identify protein and DNA/RNA structures visually in density maps. Also, we noted that the segment-based accuracy of α helices and β strands were very high, 0.950 and 0.940 for 6 Å maps, respectively (**Supplementary Table S2**). These results strongly indicate that the structure assignments by Emap2sec+ will be able to help main-chain tracing and domain structure assignments in cryo-EM maps.

Results for the 10 Å maps were about 6% to 13% worse than 6 Å maps (**Fig. 2a**). Among the four structural classes, α helices have the smallest difference, 7.5% in terms of the voxel-based F1 score and 5.6% for Q4, between 6 Å and 10 Å maps. Despite the drop, it is remarkable that the overall voxel-based F1 score and Q4 for 10 Å were maintained as high as 0.8. Therefore, we can conclude that there is rich structural information in even 10 Å simulated maps. In **Figure 2b**, we compared Q4 from the phase 1 network and the final assignments from the phase 2 network for all 108 maps in the testing set. As shown, overall Q4 values improved for almost all the maps (106 maps, 98.1%) for both 6 and 10 Å resolutions with a margin of 4.6% points and 1.5% points, respectively.

Among the four structural classes, assignments of other structures improved for the largest fraction of maps (93.5%) for 6 Å map dataset and α helices for 10 Å maps (85.2%). For DNA/RNA class, improvement by the phase 2 was observed for 77.7% of maps. As shown in **Supplementary Table S3**, the phase 2 network made consistent improvement for all the voxel-, residue/nucleotide-, and segment-based metrics for all the structural classes.

In **Supplementary Table S2**, we also computed assignment accuracy of another dataset of maps simulated at a randomly selected resolution between 6 to 10 Å. The accuracy to this dataset naturally turned out to be values between results of 6 Å maps and 10 Å maps.

**Figure 3** shows examples of the structure detection by Emap2sec+. The first three panels show performance on maps simulated at three different resolutions, 6 Å, 7.56 Å, and 10 Å, respectively. The first panel, **Figure 3a** is a complex of two protein chains of Aspartyl-tRNA synthase with two tRNAs. All structural classes are detected with high accuracy for this map with an overall Q4 accuracy of 0.842. α helices, β strands, other structures, and RNA were identified with Q4 of 0.882, 0.914, 0.695, and 1.00, respectively. Particularly, as shown in the figure, four α helices locating in front of this complex were accurately and distinctively detected. The next example, the LSR-DNA complex (**Fig. 3b**) is abundant in nucleotides and α helical residues. Besides these two abundant structural classes that were detected with a high Q4 of 0.976 and 0.890, respectively. The accuracy of β strands was lower, at Q4 of 0.670, probably mainly because they share only 10.3% of atoms in the complex. In **Figure 3c**, a 10 Å map for a ribosomal protein and RNA complex is shown. The large volume RNA structures in this map was well identified at Q4 of 0.877 despite of the low resolution. Secondary structures of proteins were detected at a slightly lower residue-based accuracy of 0.703. **Figure 3d** illustrates changes of structural assignment made by the phase 2 network using a 10 Å map as an example. This protein has a horseshoe fold that has many repeats of α helical structural motifs and contain only very small number of β-strand residues. The phase 1 result of the structure detection contains many small regions with incorrect β-strand detections that are scattered across the map. It is also observed that structure assignments are fragmented as shown in the regions indicated by circles. In contrast, the phase 2 network has smoothened and reduced noise in the structural assignments by considering assignments in neighboring voxels. The phase 2 modification improved the voxel-based F1 score from 0.806 to 0.858 and the Q4 drastically from 0.793 to 0.941. The last panel, **Figure 3e** is an example where Emap2sec+ did not perform well. This map, simulated at 6 Å resolution, contains a short RNA of 4 nucleotides. As shown in the right panel, due to the relatively small volume of RNA region, about the half of the RNA region was mis-detected as the other structures (green) as indicated by a Q4 of RNA of 0.5, because the RNA fragment is surrounded by loops. In this case, the phase 2 network made the detection of RNA worse by changing correct assignment of RNA to other structures, resulting in the voxel-based accuracy of nucleic acids reduced from 0.31 to 0.27.

**Figure 3.**
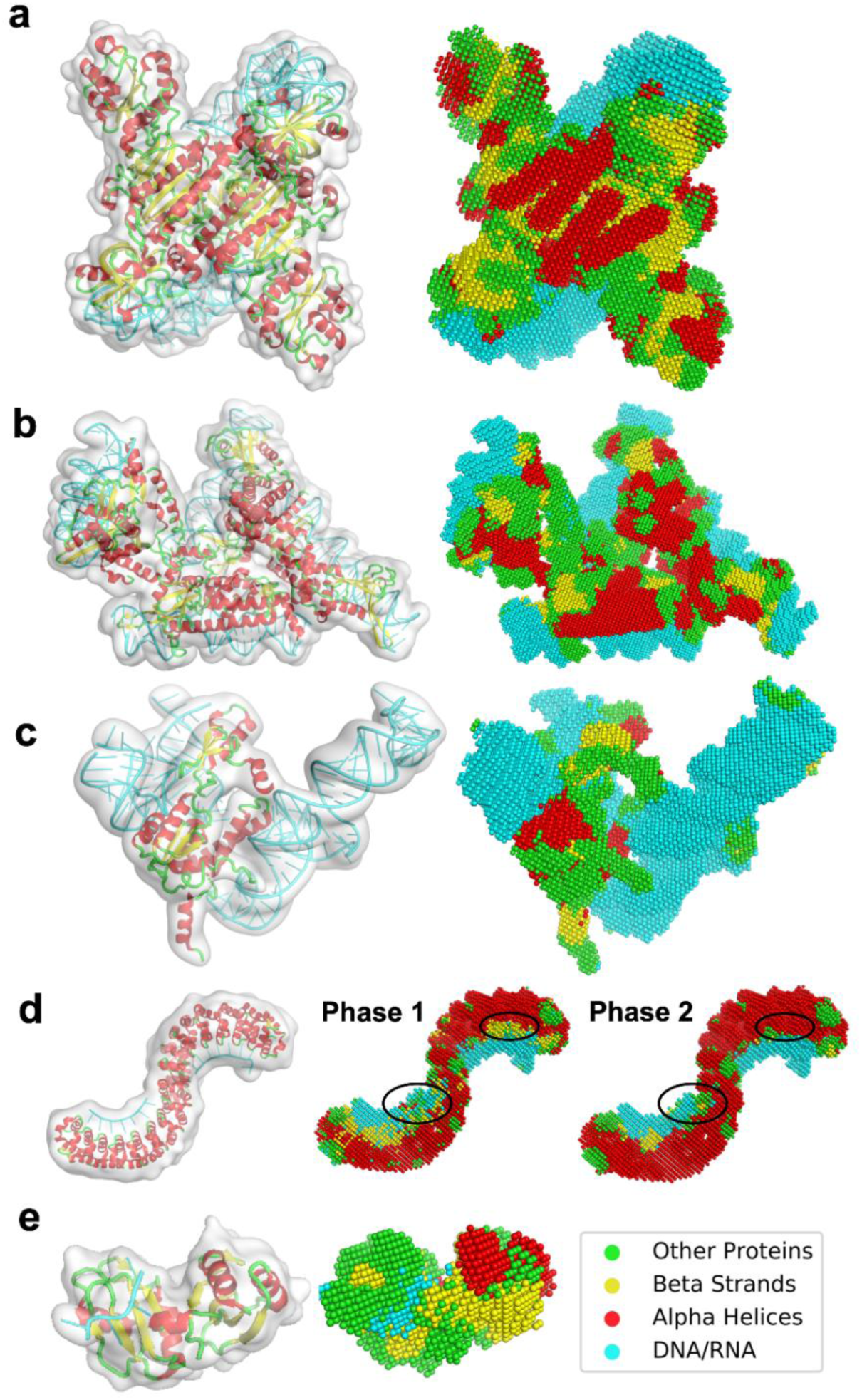
Example of the structure detection for simulated maps. For each panel, the macromolecular structure in the simulated EM map are shown on the left while the structure detection result of the phase 2 network is shown on the right. Colors of spheres in the structure detection panels indicate structure types: red α helices; yellow, β strands; green, other structures (loop); and cyan, RNA/DNA. Detailed evaluation metrics are included in Supplementary Table S3. **a**. Aspartyl-tRNA synthase complexed with tRNA(Asp) (PDB ID: 1IL2). Simulated map resolution: 6 Å. The complex contains 1170 amino acids (AA) and 129 nucleotides (nt). Voxel-based F1 score (F1): 0.879; Voxel-based accuracy (Acc): 0.880; Q4: 0.842. **b**. Large serine recombinase (LSR) – DNA complex (PDB ID: 4KIS). Simulated at 7.56 Å. 1216 AA and 208 nt. F1: 0.864; Acc: 0.863; Q4: 0.867. **c**. Ribosomal protein L30, L37a, S13 complexed with 3 ribosomal RNAs (PDB ID: 1YSH). Simulated Resolution: 10 Å. 261 AA and 163 nt. F1: 0.818; Acc: 0.800; Q4: 0.770. **d**. Pumilio homology domain complexed with RNA (PDB ID: 1M8X). Simulated resolution: 10 Å. 682 AA and 15 nt. Phase 1 results: F1: 0.806; Acc: 0.780; Q4: 0.793. Phase 2 results: F1: 0.858; Acc: 0.861; Q4: 0.941. **e**. IMP3 RRM12 in complex with RNA (PDB ID: 6GX6). Simulated resolution: 6 Å; 170 AA and 4 nt. F1: 0.712; Acc: 0.715; Q4: 0.753. Accuracies for RNA was: F1(RNA): 0.416; Acc(RNA): 0.270; Q4(RNA): 0.50.

### Structure detection on experimental maps

Next, we examined the performance of Emap2sec+ on 19 non-redundant experimental maps that are determined at a resolution between 5 to 10 Å. The networks were trained on a dataset of 83 experimental EM maps. The list of EM maps in the dataset is provided in **Supplementary Table S1**.

Q4 accuracy of individual maps are shown in **Figure 4a** relative to their map resolution. The overall Q4 (magenta) exhibits a weak negative correlation to the map resolution. More precisely, when the map resolution is better than 7 Å, more than half of the maps showed relatively high Q4 above 0.64 or higher. For the rest of the 10 maps, Q4 was stable around 0.45 to 0.5 regardless of the map resolution. It is worthwhile to note that Emap2sec+ could detect structures from the map at 10.0 Å (EMD-8131) with Q4 of 0.575, i.e. structures of about half of the residues/nucleotides were correctly identified. Q4 values of the maps are certainly lower than simulated maps that showed over a Q4 of 0.8 on average for even 10 Å maps (**Fig. 2**), indicating that structure detection is more difficult in experimental maps.

**Figure 4.**
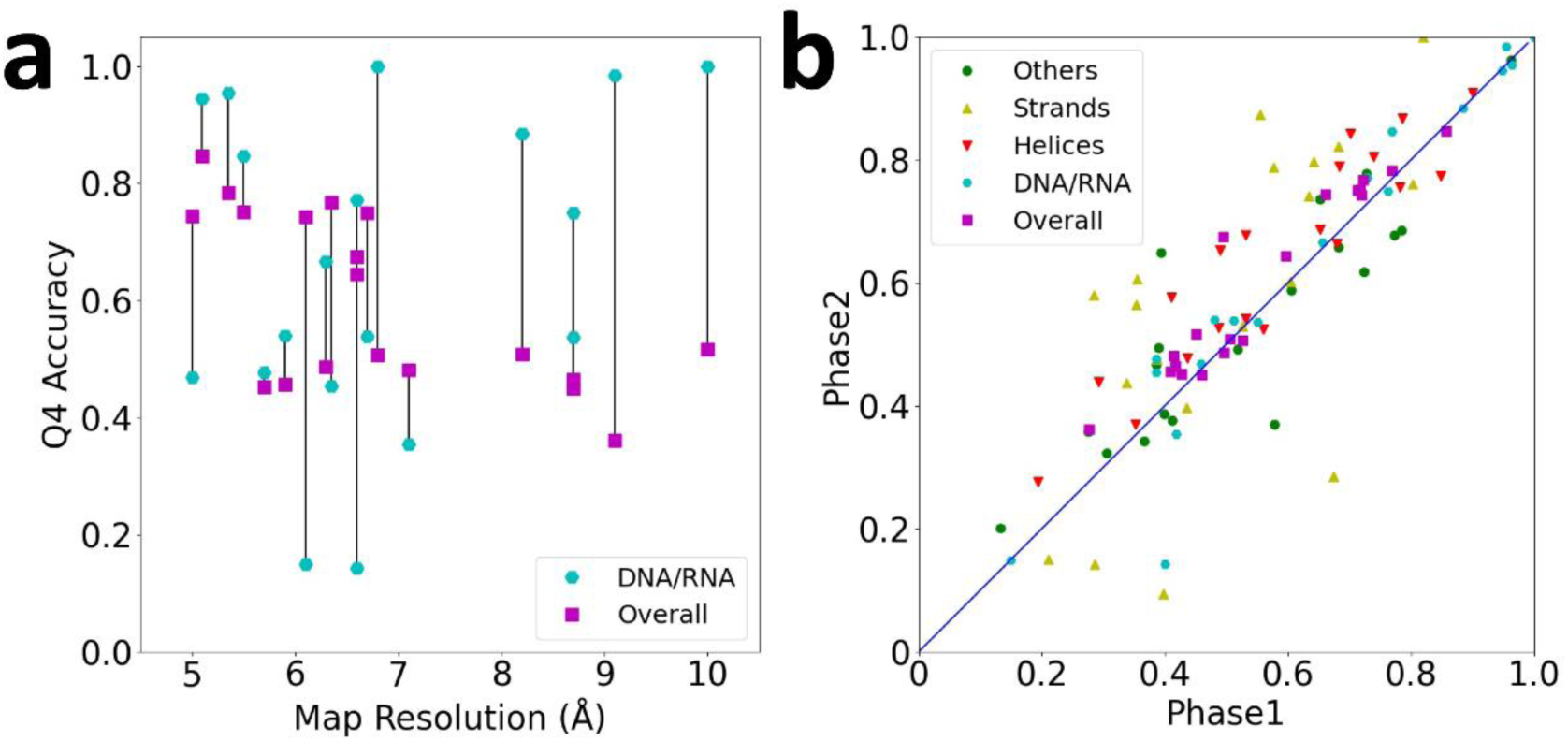
Structure class detection on 19 experimental maps. See Supplementary Table S3 for the phase 1 and phase 2 accuracy of each map. **a**. Q4 accuracy of experimental maps relative to the map resolution. Overall Q4 is shown in magenta squares and Q4 of DNA/RNA is shown in cyan circles. Lines connects values of the same map. **b**. The residue-based accuracy comparison between the Phase 1 and Phase 2 networks.

In **Figure 4a**, we also showed Q4 value of DNA/RNA (cyan circles). For 14 out of 19 maps, nucleic acids were better detected than the overall average Q4. Interestingly, detection of nucleic acids did not deteriorate as the resolution was lowered. A high Q4 for nucleic acids of over 0.8 was observed even for maps of over 8 Å resolution.

Turning our attention now to **Figure 4b**, as also observed in simulated maps, the phase 2 network improved the accuracy over the phase 1 for almost all the maps. Among the four structural classes, α helices and β strands were improved with the largest average margin of 0.058 and 0.057, respectively. Though the detection accuracy decreased for other structures for more than half of the maps, overall Q4 improved for all but four maps with an average improvement from 0.56 to 0.60.

We discuss illustrative examples of Emap2sec+’s performance on experimental maps in **Figure 5**. The first example (**Fig. 5a**) is a nucleosome where double-stranded DNA wraps around histones (EMD-3949, PDB ID: 6ESH). The map was determined at 5.1 Å. As shown in the figure, DNA was very clearly identified with a high Q4 of 0.945. α helices and other structures, which dominate histones, were also detected with a high accuracy. Q4 of α helices was 0.756, and 0.962 for other structures. As shown in the figure, α helices in histones are distinctively identified to the level that individual helix shapes are visually identified. The second map (**Fig. 5b**) is an example of an RNA-rich structure, which is the translation pre-initiation complex (EMD-4075, PDB ID: 5LMP). RNA structures are detected with a high Q4 of 0.954. Also, β strands in this map was identified well yielding 0.761 Q4 accuracy. In the zoomed window, RNA and β strand structures as well as α helices are vividly captured and β strand regions are clearly distinguished with other structures by Emap2sec+. The next map (**Fig. 5c**) is opposite to the previous one, whose volume is dominated by proteins with a DNA double-stand embedded in the middle of the structure. It is a 6.35 Å map of the dihedral oligomeric complex with four gyrase A dimers (EMD-9316, PDB ID: 6N1P). The complex holds a double-stranded DNA in the middle. Secondary structures of proteins were identified with high accuracy, particularly for α helices (Q4: 0.843), in this map. Rod shapes of a number of α helices are precisely identified. DNA structure is also detected accurately at the middle of the double strands, but the two ends were mis-recognized as loops and β strands due to lower density values than the middle part. In the last panel (**Fig. 5d**), we show the structure detection for an 8.7 Å map of TFIID-IIA complex with a promotor DNA. As shown, the DNA is clearly detected, which had a Q4 of 0.75 (only the DNA region). The overall Q4 was 0.464, which is about the average for maps with this resolution. The map did not have structure assignment for the structural region named lobe B partly due to the map resolution^22^. But interestingly, the same group later determined the structure of lobe B with a higher resolution, at 4.5 Å^23^, which is shown in the box in the second row of the panel. Compared with the newly determined structure, α helices were detected well with a high Q4 of 0.609 as visualized in the figure. With the 4.5 Å map, Emap2sec+ detected α helices with a slightly better Q4 of 0.614 and with 18% higher Q4 of 0.517 (0.438 with the 8.7 Å map) (**Supplementary Figure S4**).

**Figure 5.**
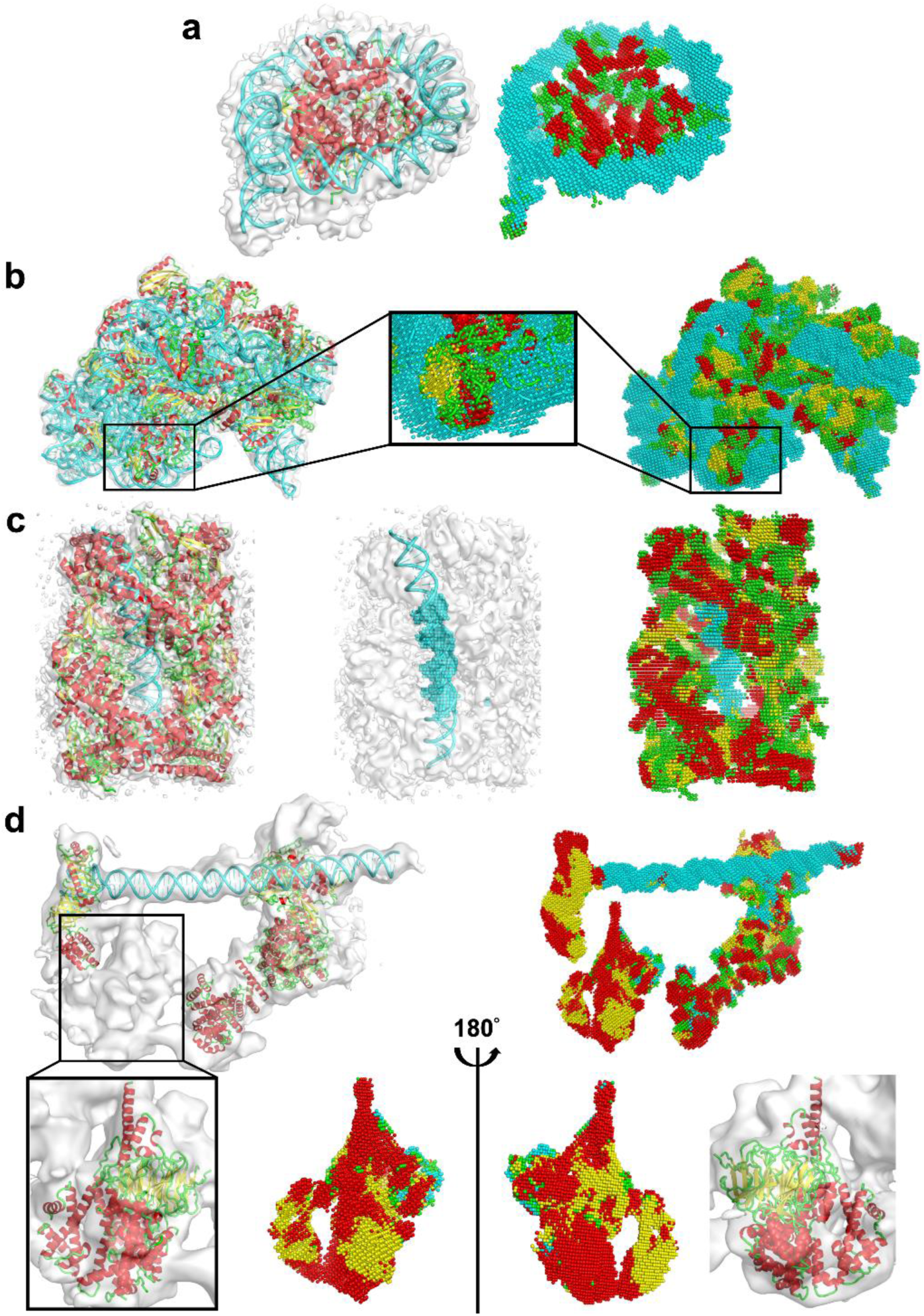
Examples of structure detection of experimental maps. The density maps and associated structures are shown on the left and the detection results of Emap2sec+ are shown on the right. Spheres in red represents detected α helices; yellow, β strands; green, other structures; and cyan, RNA/DNA. Detailed evaluation metrics are shown in Supplementary Table S4. **a**, nucleosome breathing Class 3. EMD-ID: EMD-3949; PDB ID: 6ESH. Resolution: 5.10 Å. 738 amino acids (aa) and 274 nucleotides (nt). Voxel-based F1 score: 0.887; Voxel-based accuracy (Acc): 0.870; Q4: 0.846. **b**, bacterial 30S-IF1-IF3-mRNA translation pre-initiation complex. EMD-4075; 5LMP. Res.: 5.35 Å. 2622 aa and 1534 nt. F1: 0.855; Acc: 0.846; Q4: 0.784. **c**, dihedral oligomeric complex gyrA. EMD-9316; 6N1P. Res.: 6.35 Å. 3828 aa and 88 nt. F1: 0.760; Acc: 0.749; Q4: 0.767. In the middle panel, only the DNA is shown with voxels as DNA. **d**. human TFIID-IIA bound to core promoter DNA. EMD-3305; 5FUR. Resolution: 8.7 Å; 1857 aa and 39 nt. In a box, another PDB entry, 6MZC, is shown, which is for the TFIID BC core and fills the missing structure in lobe B. 6MZC was associated with another newer EM map, EMD-9298, determined at a 4.5 Å resolution. F1: 0.487 (0.371); Acc: 0.493 (0.402); Q4: 0.516 (0.438). In the parentheses, values were shown that were computed only for the part of the structure in 6MZC that fill the density (953 aa). The structure of lob B and Emap2sec+’s detection is shown from two opposite angles. The detection results using the newer map, EMD-9298, is provided as Supplementary Figure S4.

### Comparison with related works

Prior to our work, there are very limited methods developed for detecting structures of both DNA/RNA and protein structures. Popular structure modeling tools originally developed for X-ray crystallography, ARP/wARP(v8.0)^8^, Phenix^7^, and Brickworx^24^, only work for maps that include solely proteins or DNA/RNA. The predecessor of this work, Emap2sec, is designed only for protein secondary structure detection. And as mentioned above, Emap2sec+’s performance on protein structure assignments is better than Emap2sec.

A recent work, Haruspex^19^, would be closest to Emap2sec+, as it uses a deep neural network to detect both DNA/RNA and protein structures. However, the purpose of their tool and targeted application is quite different; Haruspex is designed to detect structures in higher resolution maps at 4 Å or better to detect potential errors in structure models built from a cryo-EM or to assist modeling process while Emap2sec+ is to provide structural clue for maps at 6-10 Å where structure information is otherwise not easily detectable. Due to the difference in design, Haruspex did not work well on our low resolution datasets of simulated maps and experimental maps as shown in **Supplementary Figure S3**.

## Discussion

We reported Emap2sec+, a deep learning-based method that can detect structures in EM maps at intermediate resolution (5 to 10 Å), which substantially upgraded the previous Emap2sec by enabling detection of nucleic acids and improving protein secondary structure detection accuracy. Nucleotides were particularly well detected and the accuracy did not drop much even for maps with lower resolution. This work is the first to explore the structure information for protein-nucleic acid complexes at this difficult resolution range. The same deep learning strategy will be adopted for detecting other molecules or structures, such as amino acid types, in EM maps. Emap2sec+ will aid structural assignments and modeling with fast and accurate predictions and will be a useful and powerful tool in the era of cryo-EM structural biology.

## Methods

### Dataset of simulated and experimental EM maps

We prepared two types of datasets, simulated and experimental cryo-EM maps. The simulated EM map dataset was computed from 1,052 different PDB entries which contain protein and DNA or RNA structures. This dataset is non-redundant in that any protein pairs from two PDB entries have less than 25% sequence identity between each other. For each PDB entry, we simulated two EM maps at 6 Å and at 10 Å using the pdb2vol program of the SITUS package^25^.

We collected experimental maps containing protein and DNA/RNA structures from EMDB^3^. We first chose all cryo-EM maps that are determined at a resolution between 5 and 10 Å and have corresponding PDB entries. Maps were deleted if corresponding PDB entries have an unknown protein sequence with no amino acid type assignment. To ensure that EM maps and associated PDB structures have sufficient overlap and align properly with each other, we examined the cross-correlation of densities between the map and a simulated map from the PDB entry at the map’s resolution. Maps were removed if the cross correlation was less than 0.65. The alignment between the map and the associated PDB entry was also manually checked. Applying all these steps resulted in a dataset of 83 EM maps.

To handle the redundancy of the maps, maps that have at least one protein sequence pairs with 35% global sequence identity^26^ were clustered together using complete-linkage. This clustering resulted in 19 clusters of EM maps. We split these 19 maps into 4 folds as shown in Supplementary Table S1. For training and testing Emap2sec+, we adopted a cross-testing as follows: When each subset was used as the test set, the rest of the three subsets that were used for training were supplemented with cluster members from the 83 EM maps. This operation guarantees that there is no redundancy between the testing and the training sets; and at the same time the number of training data was enriched with the cluster members.

Density values of each map were normalized to [0.0, 1.0] with min-max normalization. For experimental maps, negative density values were set to zero before normalization and the density value of the author-recommended contour level was used as the minimum value.

The input density data for Emap2sec+ was voxels of size of 11^3^ Å^3^ collected from the simulated and experimental EM map dataset by traversing each map along the three axes with a stride of 2 Å. For each voxel, the correct structure(s) was assigned by considering the structures of heavy atoms that were within 3 Å to the center of the voxel. For protein heavy atoms, the secondary structure was assigned according to STRIDE^27^. Residues with a label of H, G, or I by STRIDE were labeled as α helix, while β strand was assigned for the labels of B/b or E. Residues with all the other labels were considered as other structures. If a heavy atom belonged to DNA or RNA, the voxel was labeled as DNA/RNA.

### Training the deep neural network of Emap2sec+

Emap2sec+ was trained separately for the set of simulated maps at 6 Å, the set at 10 Å, and the experimental maps. For simulated maps, the dataset of 6 Å or 10 Å with 1,052 simulated maps was separated into three sets, 844 maps for training and validation for the phase 1 networks, 100 maps for the phase 2 network, and the rest of 108 maps for testing. For the phase 1 networks, 844 maps were split to 80% (675 maps) and 20% (169 maps) for training and validation. The same split of 80% and 20% were applied to the 100 maps for the phase 2 network. The phase 1 networks consist of four binary classification models and a four-class classification model (**Fig. 1b**). For the binary models, an equal number of over ten million positive and negative voxels each collected from the 675 maps were used for training. Positive/negative voxels are those which have/do not have the particular structure of the binary classification. For the multi-class model, over 8.5 million voxels in total were used for training. For training the phase 2 network, we used the 100 maps, which do not have overlap with the maps used for phase 1 training and validation. After the phase 1 networks were fully trained, we input the 100 maps and obtained structure assignments to each voxels of the maps. Then, 80 maps (i.e. 80%) were used for training and 20% were used for validation. The number of training voxels for phase 2 was around 1 million.

For training the phase 1 and 2 networks, we tested learning rates of [0.00002, 0.0002, 0.002, 0.02, 0.2] using the Adam optimizer^28^ with L2 regularization with values of [1e-6, 1e-5, 1e-4, 0.001, 0.01, 0.1]. Among the combinations tested, the learning rate of 0.002 with L2 regularization parameter of 1e-5 showed the largest voxel-based accuracy on the validation set, although the differences among the combinations were small.

For training the networks for experimental maps, we used the same hyper-parameters as determined for the simulated map datasets. For evaluation four-fold cross validation was performed. Similar to the procedure applied for the networks for simulated maps, 80% of the maps in the training set were used for training the phase1 networks and the remaining 20% of the maps were used for training the phase 2 network.

### Evaluation metrics

We evaluated the performance of Emap2sec+ at three levels: the voxel, the amino acid residue/nucleotide, and the protein secondary structure segment levels. The voxel level considers if the assigned structure by Emap2sec+ agrees with the structure(s) within a 3 Å radius to the voxel center. We also evaluated with multiple structure labels for a voxel if the structures were observed within the radius. For the voxel-level evaluation, we used the F1-score, which is computed from the precision and the recall of the assignments given to the entire map or to each secondary structure class:

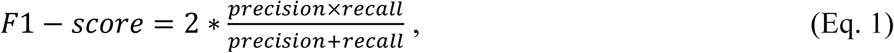

where precision is the fraction of voxels with correct structure detection over all the voxels with the structure assignment by Emap2sec+ and recall is the fraction of the voxels with correct structure detection over all the voxels that belong to the structure class. The overall accuracy (recall) and F1-score of maps were computed as the weighted average of different classes. For the residue-level evaluation, we defined Q4 accuracy, which calculates the recall for residues in each secondary structure class or nucleotides. For each residue or nucleotide, labels assigned to voxels within 3.0 Å to any heavy atoms of the residue/nucleotide were considered and the assignment was considered as correct if the majority vote from the voxels agreed with the correct class of the residue/nucleotide. For the protein secondary structures, we further reported the segment-level accuracy. A segment was defined as consecutive amino acids with same secondary structure type with the minimum length of 6 amino acids for α helix and 3 residues for a β strand. If at least 50% of the residues in a segment were assigned with the correct label, the assignment to the segment was considered as correct.

## Acknowledgments

The authors acknowledge Charles Christoffer for his help in finalizing the manuscript. This work was partly supported by the National Institutes of Health (R01GM123055), the National Science Foundation (DMS1614777, CMMI1825941, MCB1925643, DBI2003635), and the Purdue Institute of Drug Discovery. EA is supported by a fellowship from Umm Al-Qura University, Saudi Arabia.

## Author Contributions

D.K. conceived the study. X.W designed the Emap2sec+ framework with D.K., E.A., and T.W.A and X.W implemented it. The datasets were selected and analyzed by E.A. and T.W.A. A script to scan an EM map with a voxel was written by G.T. and a script for visualizing detected structures was implemented by S.R.M.V.S. The experiments were designed by X.W., E.A, T.W.A. and D.K. and were carried out by X.W. X.W. and E.A, S.R.M.V.S., G.T., and D.K. analyzed the results. The manuscript was drafted by X.W. D.K. administrated the project and edited the manuscript.

## Supplementary Information

**Supplementary Table S1.** The dataset of experimental maps.

| **Fold** | **Representative entries** | **Entries in the same clusters as representatives** |
| --- | --- | --- |
| **1** | EMD-7845 (6DBL), EMD-8518 (5U8S), EMD-6906 (5ZAM), EMD-7290 (6BU9), EMD-4671 (6QY3) | EMD-4670 (6QXT) |
| **2** | EMD-9316 (6N1P), EMD-4141 (5M1S), EMD-3545 (5MQF), EMD-0244 (6HMS), EMD-4139 (5M0R) | EMD-3198 (5FKV), EMD-3202 (5FKW), EMD-3683 (5NRL), EMD-6889 (5Z56), EMD-6890 (5Z57), EMD-9621 (6AH0) |
| **3** | EMD-6803 (5Y3R), EMD-4242 (6FEC), EMD-2810 (4D5Y), EMD-7103 (6BJS) | EMD-5942 (3J6X), EMD-5943 (3J6Y), EMD-5976 (3J77), EMD-5977 (3J78), EMD-2683 (4UJC), EMD-2682 (4UJD), EMD-2620 (4UJE), EMD-5036 (4V69), EMD-1849 (4V6K), EMD-5775 (4V7B), EMD-5799 (4V7C), EMD-5800 (4V7D), EMD-8190 (5K0Y), EMD-4078 (5LMS), EMD-4122 (5LZB), EMD-3581 (5MYJ), EMD-3661 (5NO2), EMD-3662 (5NO3), EMD-3663 (5NO4), EMD-8621 (5UZ4), EMD-7014 (6AWB), EMD-7015 (6AWC), EMD-7016 (6AWD), EMD-3637 (6FXC), EMD-0057 (6GSM), EMD-0058 (6GSN), EMD-0139 (6H58), EMD-0195 (6HCM), EMD-0197 (6HCQ), EMD-4427 (6I7O), EMD-2813 (4D67), EMD-3561 (5MS0), EMD-3696 (5NSS), EMD-7439 (6CA0), EMD-3580 (5MY1) |
| **4** | EMD-2784 (4V1M), EMD-3949 (6ESH), EMD-4075 (5LMP), EMD-3305 (5FUR), EMD-8131 (5IVW) | EMD-2785 (4V1N), EMD-2786 (4V1O), EMD-3383 (5FZ5), EMD-8132 (5IY7), EMD-8133 (5IY8), EMD-8134 (5IY9), EMD-8135 (5IYA), EMD-8735 (5VVR), EMD-8737 (5VVS), EMD-6980 (6A5L), EMD-6981 (6A5O), EMD-6982 (6A5P), EMD-6983 (6A5R), EMD-6984 (6A5T), EMD-6985 (6A5U), EMD-4182 (6F42), EMD-6986 (6INQ), EMD-0671 (6J4W), EMD-3948 (6ESG), EMD-3950 (6ESI), EMD-4336 (6G0L), EMD-9843 (6JM9), EMD-0090 (6GYK) |

EMDB ID and PDB ID are paired. Representative entries that were used as a test set in the four-fold cross-validation were shown in the Representative column. The list on the right column is entries that are similar (at least one chain in the complex has not less than 25% sequence identity to a chain in a representative entry) to one of the representative entries. Entries in the similar clusters were used for training when a different fold was used for testing. See Methods for more details of the dataset construction.

**Supplementary Figure S1.**
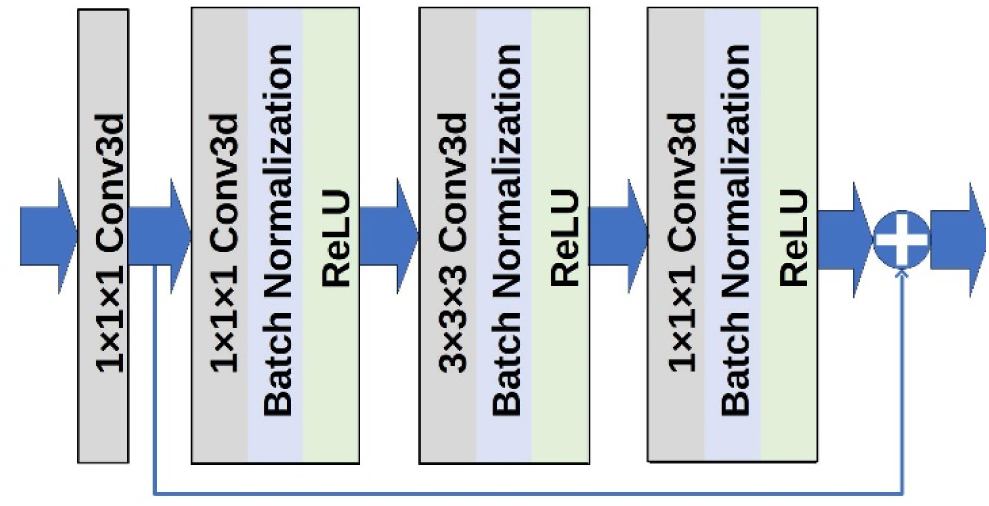
The architecture of Residual block. 6 blocks are used in the phase 1 network (Figure 1). A Residual block consists of a combination of convolutional layer, batch normalization layer, and ReLU activation with residual connections to avoid gradient collapse and previous layer’s information loss

**Supplementary Figure S2.**
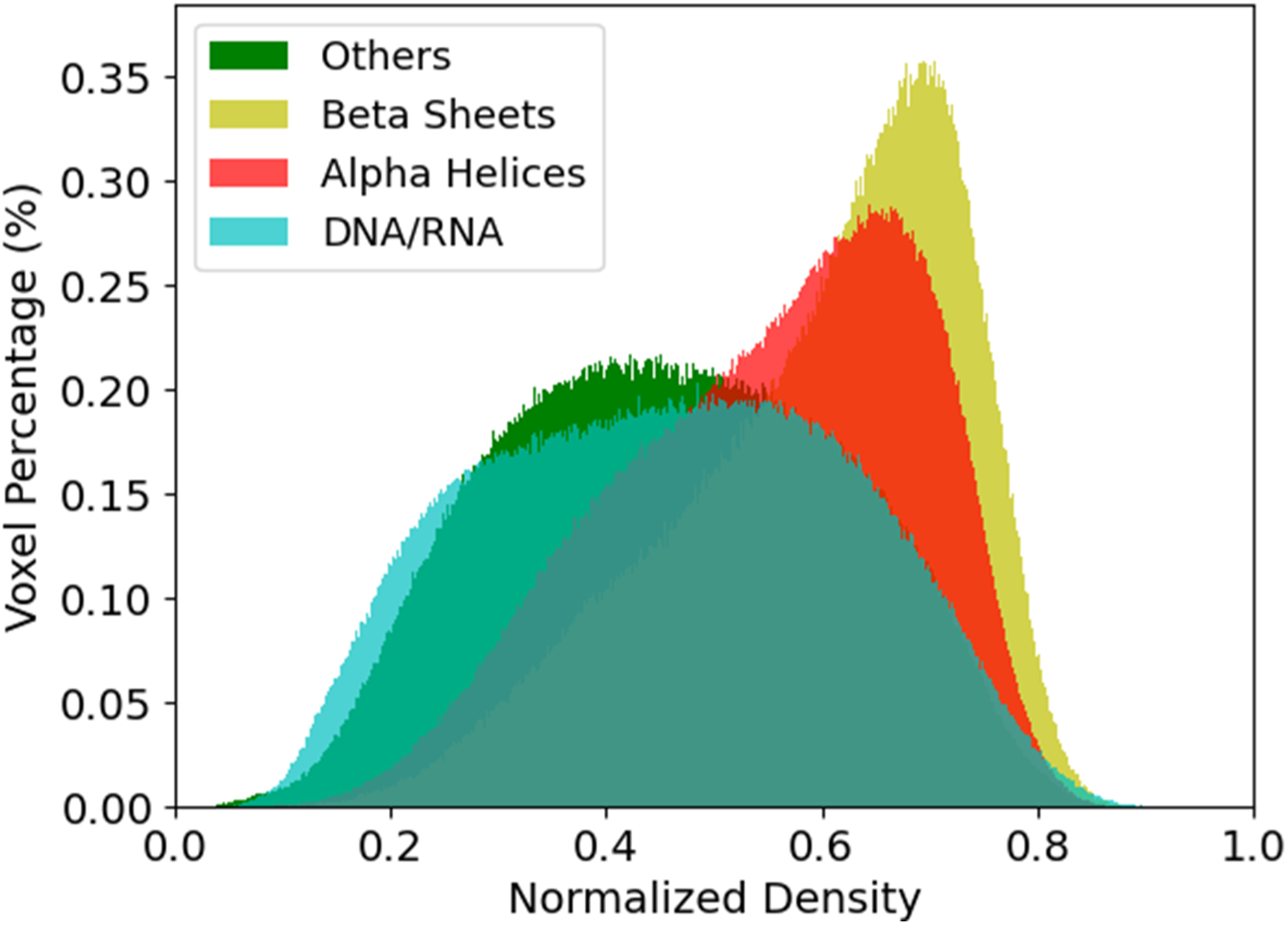
Distribution of normalized densities of four structural classes in the 108 test maps simulated at 10 Å. Corresponding distributions of 108 maps simulated at 6 Å and experimental maps are shown as Figure 2c.

**Supplementary Table S2.** Summary of the structure detection for simulated maps.

| Measure | Resolution | $\alpha$ helices | $\beta$ strands | Others | DNA/RNA | All |
| --- | --- | --- | --- | --- | --- | --- |
| **Voxel-based F1 score** | 6 Å | 0.835<br>(0.868) | 0.706<br>(0.762) | 0.723<br>(0.775) | 0.902<br>(0.907) | 0.844<br>(0.875) |
|  | 6-10 Å | 0.797<br>(0.828) | 0.640<br>(0.698) | 0.669<br>(0.719) | 0.807<br>(0.811) | 0.798<br>(0.829) |
|  | 10 Å | 0.771<br>(0.803) | 0.602<br>(0.665) | 0.634<br>(0.689) | 0.797<br>(0.801) | 0.776<br>(0.809) |
| **Voxel-based Accuracy** | 6 Å | 0.830<br>(0.857) | 0.771<br>(0.816) | 0.705<br>(0.765) | 0.907<br>(0.912) | 0.845<br>(0.876) |
|  | 6-10 Å | 0.807<br>(0.829) | 0.705<br>(0.753) | 0.650<br>(0.706) | 0.814<br>(0.817) | 0.799<br>(0.829) |
|  | 10 Å | 0.773<br>(0.796) | 0.701<br>(0.755) | 0.598<br>(0.662) | 0.805<br>(0.810) | 0.778<br>(0.810) |
| **Residue Q4** | 6 Å | 0.863<br>(0.869) | 0.816<br>(0.833) | 0.738<br>(0.767) | 0.940<br>(0.944) | 0.849<br>(0.864) |
|  | 6-10 Å | 0.838<br>(0.846) | 0.747<br>(0.775) | 0.677<br>(0.704) | 0.839<br>(0.840) | 0.804<br>(0.821) |
|  | 10 Å | 0.807<br>(0.820) | 0.751<br>(0.783) | 0.625<br>(0.666) | 0.829<br>(0.836) | 0.778<br>(0.802) |
| **Segments** | 6 Å | 0.947<br>(0.950) | 0.935<br>(0.940) | - | - | - |
|  | 6-10 Å | 0.933<br>(0.944) | 0.864<br>(0.864) | - | - | - |
|  | 10 Å | 0.904<br>(0.926) | 0.835<br>(0.892) | - | - | - |

In parentheses for different evaluation metrics, we used relaxed measure to calculate, where a cube with multiple labels will be considered as correct if model predicts one of the multiple labels. The 6-10 Å dataset contains simulated maps computed for a randomly selected resolution between 6 to 10 Å. Similar to the 6 Å and 10 Å datasets, networks for 6-10 Å were trained specifically for this dataset.
The residue Q4 is the residue/nucleoside level accuracy (recall). The segment-based accuracy considers the fraction of segments that are successfully identified. More detailed definitions are included in the Methods section.

**Supplementary Table S3. Structure assignment results for individual maps in the simulated map test set** (in a separate Excel file). Data are shown for phase 1 and 2 results for simulated maps at 6 Å, 10 Å, and at a resolution randomly selected between 6 to 10 Å. −1 in the table indicates that the structural class does not exist in the protein complex.

**Supplementary Table S4. Structure assignment results for individual maps in the experimental map test set**. (in a separate Excel file). Data are shown for phase 1 and 2 results for individual experimental maps. Fold information corresponds to the four-fold grouping of maps shown in Supplementary Table S1.

**Supplementary Figure S3.**
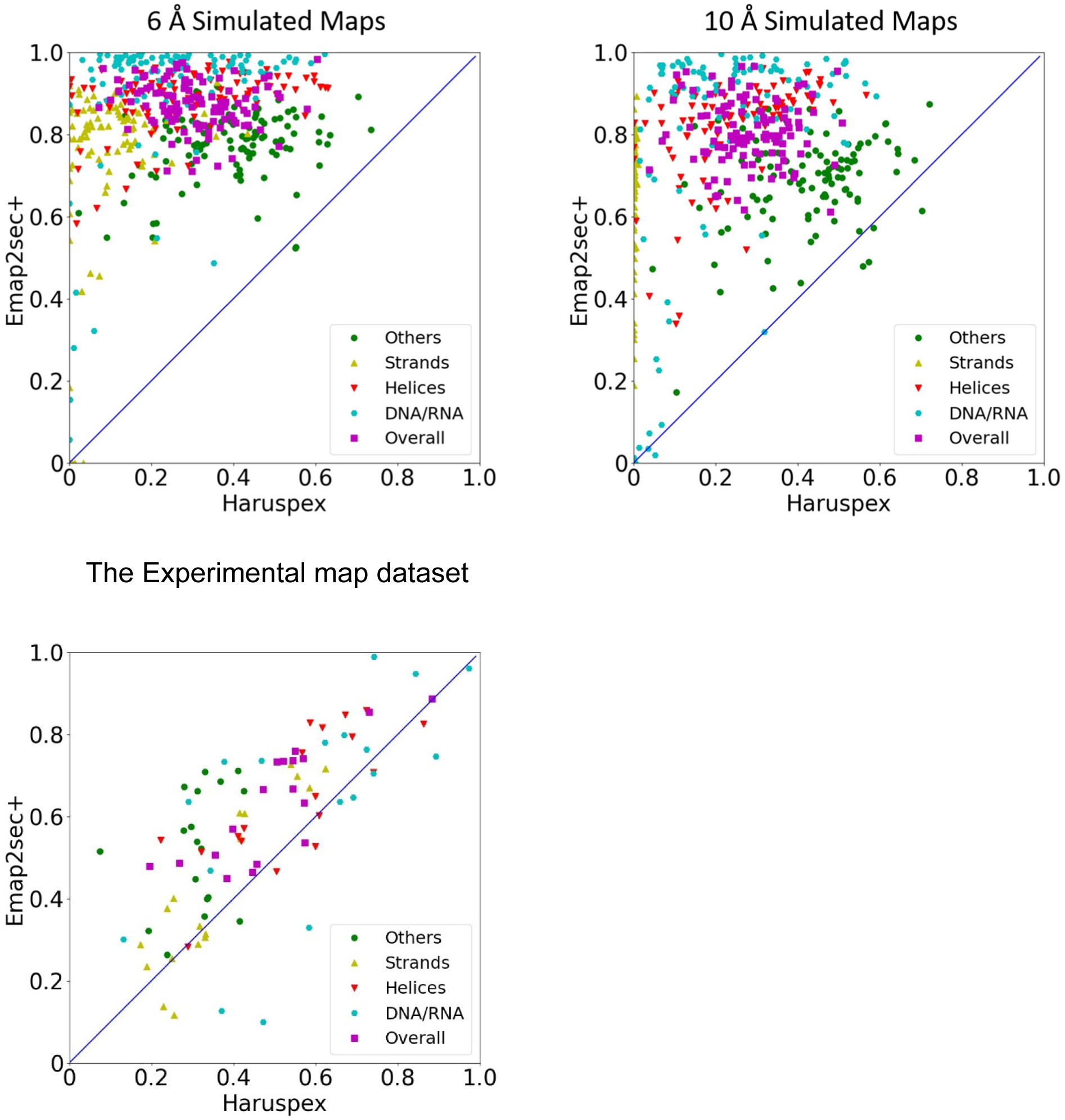
Structure detection results by Haruspex on the simulated map and the experimental map datasets. We ran Haruspex (Mostosi et al., Angew Chem Int Ed Engl, 2020) on the 6 Å and 10 Å simulated map and the experimental map datasets and compared the voxel-based F1 score with Emap2sec+. For the simulated maps at 10 Å, Haruspex detected almost no β strands.

**Supplementary Figure S4.**
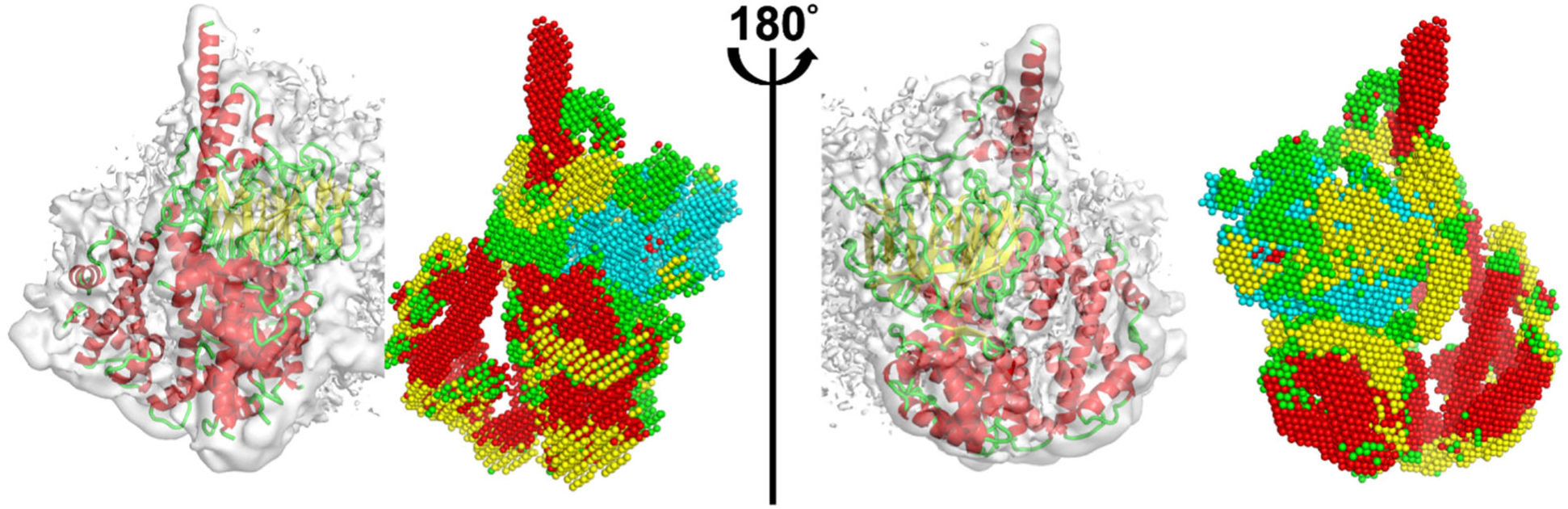
Structure detection of lob B of the TFIID-IIA and promoter DNA complex using the 4.5 Å resolution map (EMD-9298). Structures in lobe B was detected in EDM-9298, determined at 4.5 Å resolution. Detected structures were compared with corresponding region, 953 amino acids, in 6MZC. We used a contour level of 0.01. Voxel-based F1 score: 0.574; Voxel-based accuracy: 0.509; Q3(Q4): 0.517. Results for other metrics are listed in the bottom of Supplementary Table S4.

